# A bootstrap approach is a superior statistical method for the comparison of cell-to-cell movement data

**DOI:** 10.1101/2020.09.11.292672

**Authors:** Matthew G. Johnston, Christine Faulkner

## Abstract

Plasmodesmata are an increasing focus of plant research, and plant physiologists frequently aim to understand the dynamics of intercellular movement and plasmodesmal function. For this, experiments that measure the spread of GFP between cells are commonly performed to indicate whether plasmodesmata are more open or closed in different conditions or in different genotypes.

We propose cell-to-cell movement data sets are better analysed by a bootstrap method that tests the null hypothesis that means (or medians) are the same between two conditions, instead of the commonly used Mann-Whitney-Wilcoxon test. We found that that with hypothetical distributions similar to cell-to-cell movement data, the Mann-Whitney-Wilcoxon produces a false positive rate of 17% while the bootstrap method maintains a false positive at the set rate of 5% under the same circumstances. Here we present this finding, as well as our rationale, an explanation of the bootstrap method and an R script for easy use. We have further demonstrated its use on published datasets from independent laboratories.

## Main Text

Symplastic cell-to-cell connectivity is dynamically regulated in plants as a component of developmental and environmental responses (Perbal *et al*., 1996; Wada *et al*., 2002; Faulkner *et al*., 2013). Connectivity is established between cells by plasmodesmata, which function as a key parameter to define the dynamics of cell-to-cell connectivity. It is critical to assay the degree of movement of different molecules between cells to understand the range and dynamics of cell-to-cell communication as well as to assay plasmodesmal function under different conditions or in different genotypes. Accurate experimental analysis is critical to understanding this important component of plant physiology.

There are two routinely used methods, with a cellular resolution, to assay the spread of GREEN FLUORESCENT PROTEIN (GFP) from one cell into neighbouring cells: microprojectile bombardment, and low OD_600_ *Agrobacterium tumefaciens* infiltration (Oparka *et al*., 1999; Burch-Smith & Zambryski, 2010). These assays allow the experimenter to count the number of cells (‘cell count’), or the number of concentric rings of cells (‘cell layers’), to which GFP has spread from a single cell (Fig. 1a). This serves as a measure of symplastic connectivity – the further the GFP has spread, the greater the degree of connection (or of plasmodesmata permeability) between cells. Neither cell nor layer counts are parametrically distributed (Fig. **1b – d**, upper), so most studies use the non-parametric Mann–Whitney–Wilcoxon (MWW) test to compare conditions to identify factors that regulate the connection and communication between cells.

**Figure 1.**
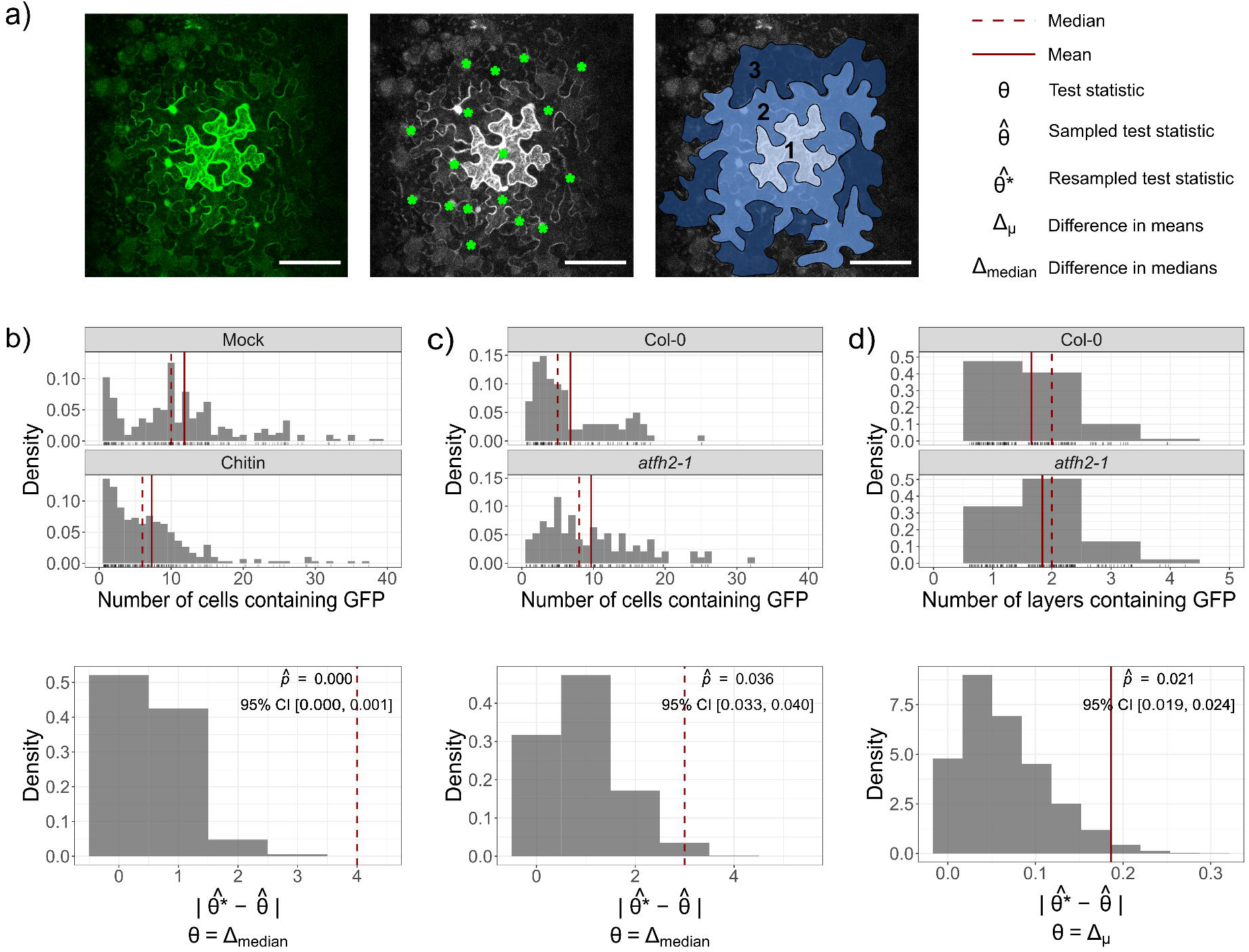
Bootstrap statistics on GFP movement data. **(a)** An example image of GFP moving from a single transformation site. The degree of movement can either be counted as the number of fluorescent cells (denoted with stars, 17 cells) or the number of cell layers with GFP (blue overlays, 3 layers). Scale bar = 100 µm. **(b – d)** *Top:* Histogram of cell counts or layers, with the median and mean marked. Bottom: Bootstrap null distributions 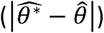 for the differences in **(b, c)** median or **(d)** mean, with estimated 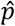 value and 95% confidence intervals (CI). The observed difference 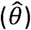 is marked by a red line. Data for **(b)** from Cheval *et al*. (2020) and data for **(c, d)** from Diao *et al*. (2018) both under the CC BY 4.0 licence.

Most experiments aim to assess whether connectivity is greater or less under different conditions, or whether plasmodesmata are more open or closed, which involves analysis of changes in the median or mean. The MWW tests the null hypothesis that two data distributions are the same (Mann & Whitney, 1947), not whether the two distributions have the same median. Therefore, it is possible to find a significant difference in an MWW test with distributions that have the same median, but different variances (Hart, 2001). When data from cell count assays are presented in histogram form, it is clear that the shape of the distributions differs between experimentally compared conditions or genotypes (Fig. **1b, c**) (Guseman *et al*., 2010; Diao *et al*., 2018; Cheval *et al*., 2020). Thus, if an MWW test is used on cell count data, the difference in distribution shapes between conditions may lead to the erroneous conclusion that there is a significant difference in the amount of spread of GFP.

Therefore, a different statistical method is required to properly interpret differences in GFP spread. For this, we propose a bootstrap method (Efron, 1979). Unlike the MWW test, bootstrapping works with data that is both non-parametric and heteroskedastic (differing variance between conditions).

The goal of the analysis is to estimate the probability that the observed difference in medians 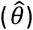 came about by chance (a *p* value). Frequentist statistics does this by comparing 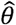 to a null distribution. In this case, the null distribution describes the probability of observing a difference in medians, when there is no true difference in the underlying data. Usually, a known distribution is used (e.g. t-distribution or F-distribution) but in this case it is unknown because the data do not follow parametric distributions (Fig. **1b, c**).

Bootstrapping techniques can be used to generate a null distribution *de novo* from the observed data already collected, as long as the samples are independent. This removes the requirement of using a known distribution. To do this, the observed data are sampled with replacement to generate a resample. This mimics what the experimenter has done originally when observing the true population. The relationship between multiple resamples and the observed data can be used to reveal how the observed data relate to the true population, and so estimate a *p* value for the observation.

An example R function is provided to perform this analysis (*medianBootstrap*.*R*, https://github.com/faulknerfalcons/Johnston-2020-Bootstrap), which requires two arguments, i.e. two vectors of numbers: control and treatment. The function generates a null distribution to compare against by resampling each vector *N* times (by default 5000) and, for each resample, generating a resampled test statistic 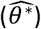. These *N* resampled test statistics are made into a null distribution by 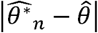 (Fig. **1b – c**, lower) as suggested by Hall and Wilson (1991).

As this is a random sampling technique, an exact *p* value cannot be calculated but an estimate is produced: a Monte Carlo 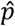 value (Eqn. 1). To do so, 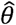 is compared to the null distribution to find the chance of observing a value at least as extreme (line on Fig. **1b – c**, lower). A +1 is added to the numerator and denominator in Eqn. 1 as suggested by Davison and Hinkley (1997): conceptually, this can be considered as including the observed sample among the bootstrap resamples.

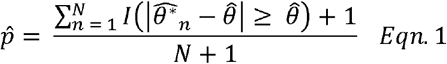

where *I* (·) is the indicator function.

As 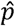 is an estimate of *p*, a 95% confidence interval should be constructed, where *p* will fall within this range 95% of the time (Wilson, 1927).

This method is not confounded by differences in variance or shape as with the MWW test. To illustrate this, we compared the Type I error rate (false positives) between the MWW and *medianBootstrap tests*, when testing if there is a difference in medians between two populations for which there was no true difference in medians, i.e. *θ* = 0. In this scenario, an error rate of 5% is expected at *α* = 0.05. Samples (*n* = 100) for each population were drawn from normal distributions with the same variance (*X, Y* ∼*N* (0,1)) simulated in R 4.0.0 (R Core Team, 2020). Both the MWW and *medianBootstrap* methods gave a difference in medians about 5% of the time, as expected (4.5% (95% CI [3.4, 6.0]) and 4.9% (95% CI [3.7, 6.4]), respectively). When variances differed between populations (*X*∼*N* (0,1), *Y* ∼*N* (0, 5^2^)), the MWW test had a false positive rate significantly higher than the set 5% of 7.5% (95% CI [6.0, 9.3]). Conversely, the false positive rate of the *medianBootstrap* method was correctly controlled at 4.7% (95% CI [3.6, 6.2]).

Alternatively, when two samples are drawn from populations with equal variance and median, but differing shape 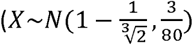, *Y* ∼ *Beta* (1,3)), a *medianBootstrap* method finds a significant difference in 5.1% of the trials (95% CI [3.9, 6.6]), as expected. Whereas, an MWW test inflates the Type I error rate to 17% (95% CI [15, 19]). Therefore, as cell count data exhibit unequal variances and differing distribution shapes between conditions and/or genotypes, we propose that bootstrap methods are a more appropriate analysis to identify differences in the spread of GFP. It is worth noting that any test statistic, *θ*, can be computed in a bootstrapped manner, provided the test is invariant to scaling. This means bootstrap testing can be extended to cell layer data, where means should be compared, as there is no difference in medians (Fig. **1d**). An example of this extension is given in *medianBootstrap*.*html*.

We acknowledge alternative advanced statistical techniques, such as linear mixed effects models, for the analysis of these data. However, they require more assumptions and are less user-friendly. We consider this bootstrap method a good, easy-to-use, superior alternative to MWW analysis of cell-to-cell movement data.

## Acknowledgments

We thank Joanna Jennings (Department of Crop Genetics, John Innes Centre) for providing the confocal micrograph in Fig. **1a** and Dr Joshua Hodgson (Department of Medicine, University of Cambridge) and Dr Matthew Castle (Department of Genetics, University of Cambridge) for constructive comments on the manuscript. The data in Fig. **1** comes from **(b)** Figure S2 of Cheval et al. (2020), **(c)** Figure 2d Diao *et al*. (2018), **(d)** Figure 2c Diao *et al*. (2018) under use of the CC BY 4.0 licence. MGJ is funded by a John Innes Foundation Studentship. Research in the Faulkner lab is supported by the Biotechnology and Biological Research Council Grant (BB/L000466/1, BBS/E/J/000PR9796) and the European Research Council (725459, “INTERCELLAR”).

## Author Contributions

MGJ and CF designed, discussed and wrote up the research. MGJ performed the analysis.

